# Addressing the mean-variance relationship in spatially resolved transcriptomics data with *spoon*

**DOI:** 10.1101/2024.11.04.621867

**Authors:** Kinnary Shah, Boyi Guo, Stephanie C. Hicks

## Abstract

An important task in the analysis of spatially resolved transcriptomics data is to identify spatially variable genes (SVGs), or genes that vary in a 2D space. Current approaches rank SVGs based on either *p*-values or an effect size, such as the proportion of spatial variance. However, previous work in the analysis of RNA-sequencing identified a technical bias, referred to as the “mean-variance relationship”, where highly expressed genes are more likely to have a higher variance. Here, we demonstrate the mean-variance relationship in spatial transcriptomics data. Furthermore, we propose *spoon*, a statistical framework using Empirical Bayes techniques to remove this bias, leading to more accurate prioritization of SVGs. We demonstrate the performance of *spoon* in both simulated and real spatial transcriptomics data. A software implementation of our method is available at https://bioconductor.org/packages/spoon.

## 1 Introduction

Advances in transcriptomics have led to profiling gene expression in a 2D space using spatially resolved transcriptomics (SRT) technologies [1]. These technologies have already led to novel biological insights across diverse application areas, including cancer [2], developmental biology [3, 4], and neurodegenerative disease [5, 6]. These emerging data types have also motivated new computational challenges, such as spatially-aware quality control to identify low-quality observations [7] and spatially-aware clustering to identify discrete spatial domains [8]. Another common data analysis task with these data is to perform feature selection by identifying a set of spatially variable genes (SVGs) [9–14]. The top SVGs are identified by ranking the genes based on some metric, such as *p*-values or an effect size like the proportion of spatial variance [10]. Accurately identifying SVGs is important because the top features are often used for downstream analyses, such as dimensionality reduction or unsupervised clustering [15–19].

Recently, Weber et al. [9] developed a computational method to identify SVGs based on a nearest-neighbor Gaussian process (NNGP) regression model [20]. In the paper, the authors identified an important relationship in SRT data. Specifically, they found a relationship between the estimated spatial variation and the overall expression, where genes that have higher overall expression are more likely to be more spatially variable. This phenomenon, known as the “mean-variance relationship”, is a well-documented technical bias in genomics [21–29]. As previously shown in other sequencing-based technologies, the reason for this bias is due to the preprocessing and normalization steps that are often applied to raw gene expression counts, or the number of unique molecular identifiers (UMIs) mapping to each gene. Specifically, Weber et al. [9] used normalized log_2_-transformed gene expression as input to the NNGP model. These preprocessing techniques are widely used in bulk RNA-seq, scRNA-seq, and SRT data, because these transformations are assumed to enable the use of statistical models based on Gaussian distributions, rather than less tractable count-based distributions [10, 28, 30–32].

However, previous work in the analysis of bulk and scRNA-seq data has also shown that because counts have unequal variances (or larger counts have larger standard deviations compared to smaller counts [33]) (**Figure S1A**), applying these log-transformations is problematic as it can overcorrect (or large logcounts can have a smaller standard deviation than small logcounts) (**Figure S1B**). In these settings, it is important to account for the mean-variance relationship. Another way to think about the mean-variance relationship is to describe it as heteroskedasticity [34] in the context of using linear models. In contrast, homoskedasticity, in the case of profiling gene expression, would be if all genes in a sample had the same variance. When applying statistical models that assume homoskedasticity in the data, if we ignore the mean-variance relationship, our results would produce inefficient estimators or even incorrect results [22, 35, 36]. For example, in differential expression analysis, ignoring the mean-variance relationship can produce false positive differentially expressed genes [25].

To address this technical bias in SRT data, here we introduce the *spoon* framework, which was inspired by the limma-voom method [33] developed for bulk RNA-seq data. In this way, the name *spoon* incorporates the concepts of both “spatial” and its origin in RNA-seq. Using real and simulated SRT data, we show that *spoon* is able to correct for the mean-variance relationship leading to more accurately prioritizing SVGs. A software implementation of our method is available as an R/Bioconductor package (https://bioconductor.org/packages/spoon).

## 2 Materials and Methods

### 2.1 An overview of the *spoon* model and methodological framework

The *spoon* model was inspired by the limma-voom method [33], which estimates the mean-variance relationship to obtain precision weights for each observation to be used as input into a linear regression model to identify differentially expressed genes with bulk RNA-sequencing data [26]. In *spoon*, we use a similar idea. First, we use Empirical Bayes techniques to estimate observation- and gene-level weights. However, here we use a Gaussian process regression model, rather than a linear regression model, to model SRT data. Then, we leverage the Delta method to re-scale the data and covariates by these weights to address the heteroskedasticity in SRT data. Briefly, the Gaussian process (GP) regression model is specified as follows [20]:

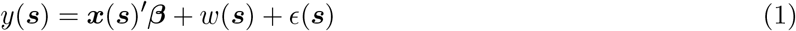

where ***s*** are the spatial locations, *y*(***s***) is the response at a location, ***x***(***s***) is a vector of explanatory variables, *w*(***s***) is a function accounting for the spatial dependence, and *ϵ*(***s***) *∼ N* (0, *τ* ^2^) is noise. ***β*** is a fixed effect, while *w* and *ϵ* are random effects. *w*(***s***) is modeled with a Gaussian process, *w*(***s***) *∼ GP* (*μ*(***s***), *C*(***θ***)), where *μ*(***s***) is a mean function and *C*(***θ***) is a covariance function with parameters *θ* = (*σ*^2^, *ϕ*, …) for the Matérn covariance function:

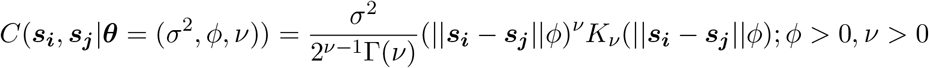

where *σ*^2^ is the spatial component of variance, *ϕ* is the decay in spatial correlation, *ν* is the smoothness parameter, and *K*_*ν*_ is the Bessel function of the second kind with order *ν*. Because we fit these models on a per-gene basis with up to thousands of genes in a given dataset, we use a nearest-neighbor Gaussian process (NNGP) [37, 38] to reduce the computational running time and make *spoon* useful to practitioners. The key idea behind using NNGPs is that instead of conditioning on all of the points in the data, only a subset (a set of nearest neighbors) of the data are used for the conditioning. Conditioning on enough of the closest neighbors provides sufficient estimates of the needed information needed and improves storage and computational costs. Briefly, a NNGP is fit to the preprocessed expression values for each gene:

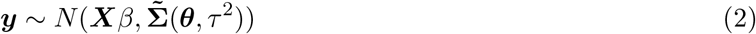

where the primary difference between a full GP model (**Equation 1**) and a NNGP (**Equation 2**) is that the NNGP covariance matrix, 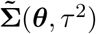, is a computationally fast approximation to the covariance matrix from a full GP model, **Σ**(***θ***, *τ* ^2^) = ***C***(***θ***) + *τ* ^2^***I***. In other words, 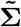 approximates the covariances from both from *w*(***s***) and *ϵ*(***s***). For the kernel, *C*(***θ***) = [*C*_*ij*_(***θ***)], we assume an exponential covariance function:

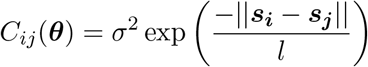

where ***θ*** = (*σ*^2^, *l*), and *σ*^2^ is the spatial component of variance of interest. *σ*^2^ is different from the nonspatial component of variance, *τ* ^2^, which is also referred to as the nugget. *l* is the lengthscale parameter, which sets how quickly the correlation decays with distance. ||***s***_***i***_ *−* ***s***_***j***_|| is the Euclidean distance between spatial locations. To estimate the parameters in the NNGP model, we use the BRISC R package [20]. Using the estimated parameters, we calculate an effect size, the proportion of spatial variance 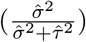.

### 2.2 Calculating observation- and gene-level weights using Empirical Bayes techniques

Briefly, we calculate the average log_2_ expression values and the standard deviations of the residuals from fitting an NNGP model per gene using BRISC (**Figure 1A**). Then, we use splines to fit the gene-wise mean-variance relationship (**Figure 1B**). Finally, we use the fitted curve to estimate observation- and gene-level weights (**Figure 1C**). Next, we describe each of these steps in greater detail.

**Figure 1:**
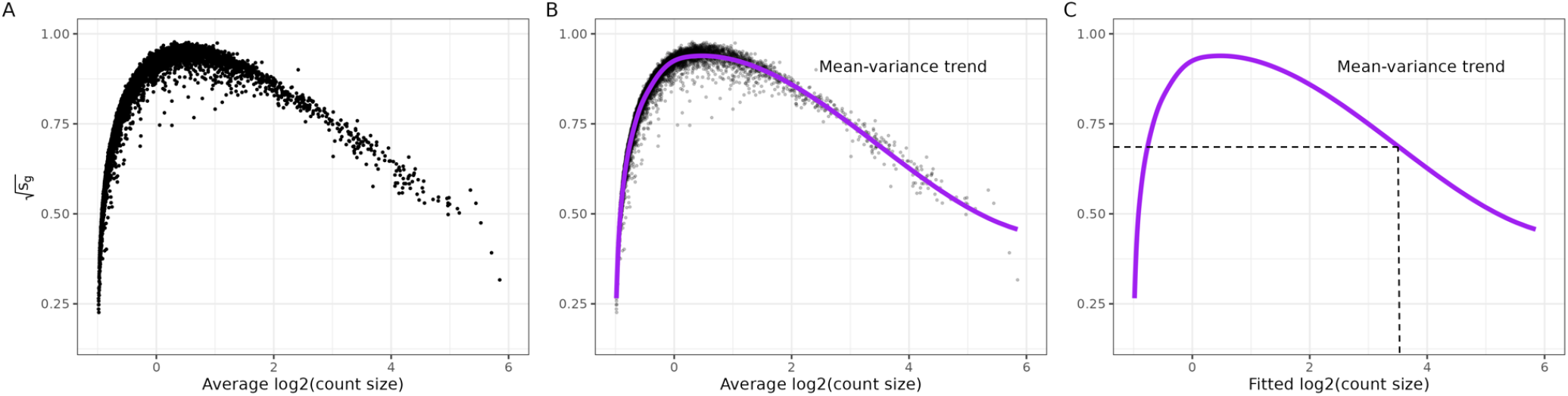
Calculating precision weights for individual observations. These data are from Invasive Ductal Carcinoma breast tissue analyzed with 10x Genomics Visium [39], hereafter referred to as “Ductal Breast”. (**A-C**) The square root of the residual standard deviations estimated using nearest neighbor Gaussian processes (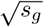 defined in Equation 3) are plotted against average logcount 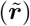. (**B**) Same as A, except a spline curve (purple) is fitted to the data to estimate the gene-wise mean-variance relationship. (**C**) Using the fitted spline curve, each predicted count value 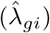 is mapped to its corresponding square root standard deviation value using 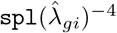.

#### 2.2.1 Fitting per-gene NNGP models using logCPM values

We start with a counts matrix, transposed so each row is a spot and each column is a gene. There are *n* spots and *G* genes in the counts matrix. The UMI counts can be indexed by *r*_*gi*_ for spots *i* = 1 to *n* and genes *g* = 1 to *G*. We define the total number of UMIs for sample *i* as 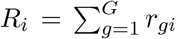. Next, we transform *r*_*gi*_ to adjust for the total number of UMIs (*R*_*i*_) by using logcounts per million (logCPM). We use a pseudocount of 0.5 to ensure we do not take the log of 0 and we add a pseudocount of 1 to the library size to make sure 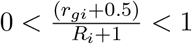:

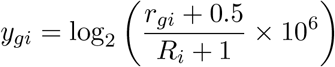

Using the normalized and log_2_-transformed data *y*_*gi*_, we fit a NNGP model (**Equation 2**) per gene with a default of ***X*** = **1**_[*N ×*1]_, corresponding to including an intercept, with *β*_*g*_ representing the overall mean expression level for gene *g*. Using the observed data *y*_*gi*_ and the predicted value 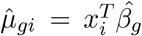, we can calculate the standard deviation of the residuals between *y*_*gi*_ and 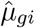:

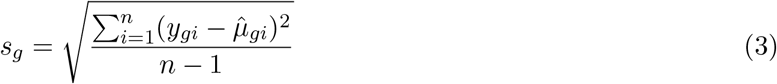

The square root of *s*_*g*_ is what we use to represent the ‘variance’ in the mean-variance relationship (see *y*-axis in **Figure 1A-C**). This concept is used in limma-voom as well because the square root of the standard deviations is roughly symmetrically distributed.

#### 2.2.2 Modeling the mean-variance relationship using 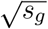 and 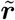

Next, we fit a nonparametric spline curve to model the mean-variance relationship in our data. Instead of using 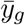 directly to represent the ‘mean’ component, we convert 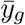 to average logcount using the geometric mean of library size, 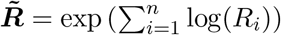. We use the geometric mean to avoid integer overflow:

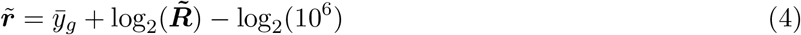

Then, we use smoothing splines (specifically `smooth.spline()` in the base R `stats` package) to model the mean-variance relationship between 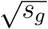 and 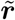. We use splines because we found they are a robust way to model the mean variance relationship seen across multiple datasets. We use the notation `spl()` to denote the fitted curve (**Figure 1B**), which represents an estimate of the mean-variance relationship.

#### 2.2.3 Prediction modeling using fitted spl() curve

Similar to Equation 4, we convert the predicted value 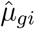 (on the logCPM scale) to a predicted count value:

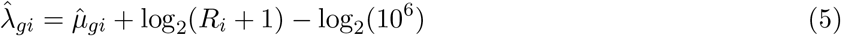

The fitted counts values for each observation are used as input to predict the square root residual standard deviation values for each *y*_*gi*_ using the spline curve. **Figure 1C** shows an example of mapping an individual observation to a square-root standard deviation value using its fitted value from the BRISC models.

To avoid extrapolating beyond the range of the function, individual observations that have 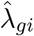 more extreme than the range of 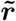 are constrained. If 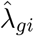 is greater than 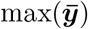, then the predicted square root residual standard deviation value for that observation is constrained to 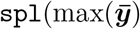). If 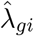 is less than 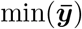, then the predicted square root residual standard deviation value for that observation is constrained to 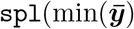). The final step is taking the inverse of the squared predicted standard deviation to compute the weight for each individual observation. The weight for each observation is defined as 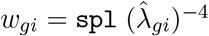, using the constrained values for observations outside of the range.

### 2.3 Correct for heteroskedasticity using observation- and gene-level precision weights

If the desired SVG detection method accepts observation- and gene-level weights, then the estimated weights *w*_*gi*_ (described in Section 2.2) can be used as input directly into the method. If the desired SVG detection method does not accept weights, then the Delta method is leveraged to rescale the data and covariates by the weights. These scaled data and covariates are used as inputs into the desired SVG detection function.

For example, the SVG detection tool called nearest neighbor SVGs (nnSVG) [9] uses a Gaussian process regression model and can have weights incorporated in the following way. We correct for the heteroskedasticity by adjusting with precision weights, *w*_*gi*_ for gene *g* at spatial location *i*. If ***W*** is a diagonal matrix where each diagonal element is *w*_*gi*_, then we know:

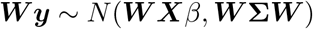

where

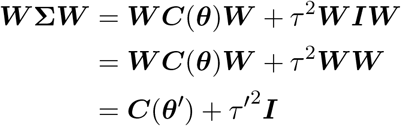

and the new input data to nnSVG would be ***W y*** and ***W X*** where 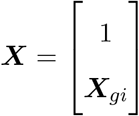

### 2.4 Data

#### 2.4.1 Real SRT data

Tissues from several regions of the human body analyzed with 10x Genomics Visium were used in the analyses. The datasets and preprocessing steps are further described below:

##### Ductal Breast

Invasive Ductal Carcinoma breast tissue data are publicly available from the 10x Genomics website. It contains one tissue sample from one donor with Invasive Ductal Carcinoma [39]. After preprocessing, this dataset contains 12,321 genes and 4,898 spots.

##### Lobular Breast

Invasive Lobular Carcinoma breast tissue data are publicly available from the 10x Genomics website. It contains one tissue sample from one donor with Invasive Lobular Carcinoma [40]. After preprocessing, this dataset contains 12,624 genes and 4,325 spots.

##### Subtype Breast

Estrogen receptor positive (ER+) breast cancer tissue data are publicly available on Zenodo and contains several tissue samples of breast cancer tissue. Only sample CID4290 is used for this analysis [41]. After preprocessing, this dataset contains 12,325 genes and 2,419 spots.

##### DLPFC

This dataset contains two pairs of spatial replicates of human postmortem dorsolateral prefrontal cortex (DLPFC) tissue from three neurotypical adult donors. Only tissue sample 151507 is used for this analysis [15]. After preprocessing, this dataset contains 7,343 genes and 4,221 spots.

##### HPC

This dataset contains human postmortem hippocampus (HPC) tissue from several neurotypical adult donors. Each sample was broken up into four Visium slides due to the large size of the HPC. Only tissue sample V12D07 335, portion D1 is used for this analysis [16]. After preprocessing, this dataset contains 5,348 genes and 4,992 spots.

##### LC

This dataset contains human postmortem locus coeruleus (LC) tissue from five neurotypical adult donors. Only tissue sample 2701 is used for this analysis [42]. After preprocessing, this dataset contains 1,331 genes and 2,809 spots.

##### Ovarian

This dataset contains tissues collected during interval debulking surgery from eight highgrade serous ovarian carcinoma patients undergoing chemotherapy. Only one tissue sample from patient 2 is used for this analysis [43]. After preprocessing, this dataset contains 12,022 genes and 1,935 spots.

Preprocessing was performed as uniformly as possible across the datasets. For datasets that had an annotation for whether or not a spot was in the tissue, spots outside of the tissue were removed. For the Subtype Breast dataset, spots that were classified as artifacts were removed. `nnSVG::filter genes()` was used to remove genes without enough data, specifically we kept genes with at least 2 counts in at least 0.2% of spots. For the LC dataset, we used a UMI filter instead of this function to remove genes with less than 80 total UMI counts summed across all spots. `scuttle::logNormCounts()` with default arguments was used to compute log-normalized expression values.

#### 2.4.2 Simulated SRT data

To simulate the mean-variance relationship, we simulated raw gene expression counts following a Poisson distribution:

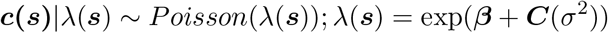

where ***s*** are spatial locations, ***β*** is a vector of true mean expression per gene, *σ*^2^ is the spatial component of variance, and ***C*** is the covariance function using a Matérn kernel with squared exponential distance. The *σ*^2^ values and ***β*** values were randomly assigned from ranges of [0.2, 1] and [ln(0.5), ln(1)], respectively. We intentionally simulate *σ*^2^ and *β* values so they are not correlated. In this way, we ensure we are simulating SVGs at all levels of mean expression. A fixed lengthscale parameter was chosen for all of the genes in a given simulation. Based on the estimated lengthscale distributions for four datasets, we chose to focus our simulations on smaller lengthscales because the majority of estimated lengthscales are between 0 to 0.15 (**Figure S2**). For reference, a scaled lengthscale value of 0.15 is interpreted as 15% of the maximum width or height of the tissue area on a standard Visium slide. We simulated 1000 genes in the following simulations.

In addition, we also considered the performance as a function of varying the lengthscale parameter *l* in *θ* = (*σ*^2^, *l*). In the NNGP model, the lengthscale parameter sets how quickly the correlation decays with distance. In the nnSVG SVG detection method [9], a key innovation was using a flexible lengthscale parameter to fit the model for each gene. Genes within the same tissue can spatially vary with different ranges of sizes and patterns, so a flexible lengthscale parameter for each gene enables the discovery of distinct biological processes. For the primary simulation evaluation, a lengthscale of 100 was used. This corresponds to a scaled lengthscale value of roughly 0.02. For supplementary simulation evaluations, 50, 60, 100, and 500 lengthscales were used. These correspond to 0.010, 0.012, 0.020, and 0.100 of the maximum width or height of the tissue area on a standard Visium slide. The spatial coordinates from the example dataset `Visium_DLPFC()` in the `STexampleData` package were used. This dataset contains 4,992 spots. We used the subset of 968 spots with row and column coordinates between 20 to 65 as the spatial coordinates to reduce the amount of time to simulate data.

### 2.5 Methods to detect SVGs

For Moran’s I [44], we ranked genes by the Moran’s I value. For nnSVG [9], the genes were ranked within the method based on the estimated likelihood ratio test statistic values comparing the fitted model against a classical linear model, assuming the spatial component of variance is zero. For SpaGFT [45], the gene ranks were calculated within the method based on decreasing GFTscore, a measure of randomness of gene expression. For SPARK-X [11], adjusted combined *p*-values from multiple covariance matrices and kernels were used to rank genes. For SpatialDE2 [46], the genes were ranked by the negative of the fraction of spatial variance for each gene. All of the criteria were ranked using the `ties.method = ‘‘first’’` option.

1. Moran’s I: `Rfast2::moranI()` [47] was used to compute Moran’s I values, and the negative Moran’s I value for each gene was ranked.
2. nnSVG: `nnSVG::nnSVG()` [9] was used, and the rank was calculated as part of the output of the function.
3. SpaGFT: `SpaGFT.detect svg()` [45] was implemented in Python, and the rank was calculated as part of the output of the function.
4. SPARK-X: `SPARK::sparkx()` [11] was run with the option of a mixture of various kernels. The combined *p*-value from all the kernels for each gene was ranked.
5. SpatialDE2: `SpatialDE.fit()` [46] was implemented in Python to fit the model for each gene. The negative of the fraction of spatial variance for each gene was ranked.

An intercept-less covariate matrix is required to implement a weighted version of an SVG detection method. To the best of our knowledge, nnSVG is the only SVG detection tool with the option to include a covariate matrix without an intercept term. The weights from *spoon* have the potential to integrate with other methods based on the flexibility of their design.

### 2.6 Code Availability

*spoon* is freely available for use as an R package available from Bioconductor at https://bioconductor.org/packages/spoon. The code to reproduce the analyses in this paper is available on GitHub at https://github.com/kinnaryshah/MeanVarBias. We used *spoon* version 1.1.3 and R version 4.4.1 for the analyses in this manuscript.

## 3 Results

### 3.1 The mean-variance relationship exists in spatial transcriptomics data

We begin by systematically demonstrating the mean-variance relationship in SRT data. This finding builds upon the initial finding suggested in Weber et al. [9]. In contrast to investigating this bias in one tissue from one tissue section, here we explore this finding across multiple tissue sections from different regions in the human body, namely DLPFC, Ductal Breast cancer, HPC, LC, and Ovarian cancer. To visualize the mean-variance relationship, we plot the mean logcounts against different components (spatial and non-spatial components) of variance calculated using `nnSVG`. As seen in **Figure 2**, the mean-variance relationship is a concern in SRT data, specifically in the nonspatial component of variance, *τ* ^2^. Given *τ* ^2^ is used when calculating the proportion of spatial variance, this suggests the way genes are prioritized as spatially variable is dependent on the overall mean expression for the gene.

**Figure 2:**
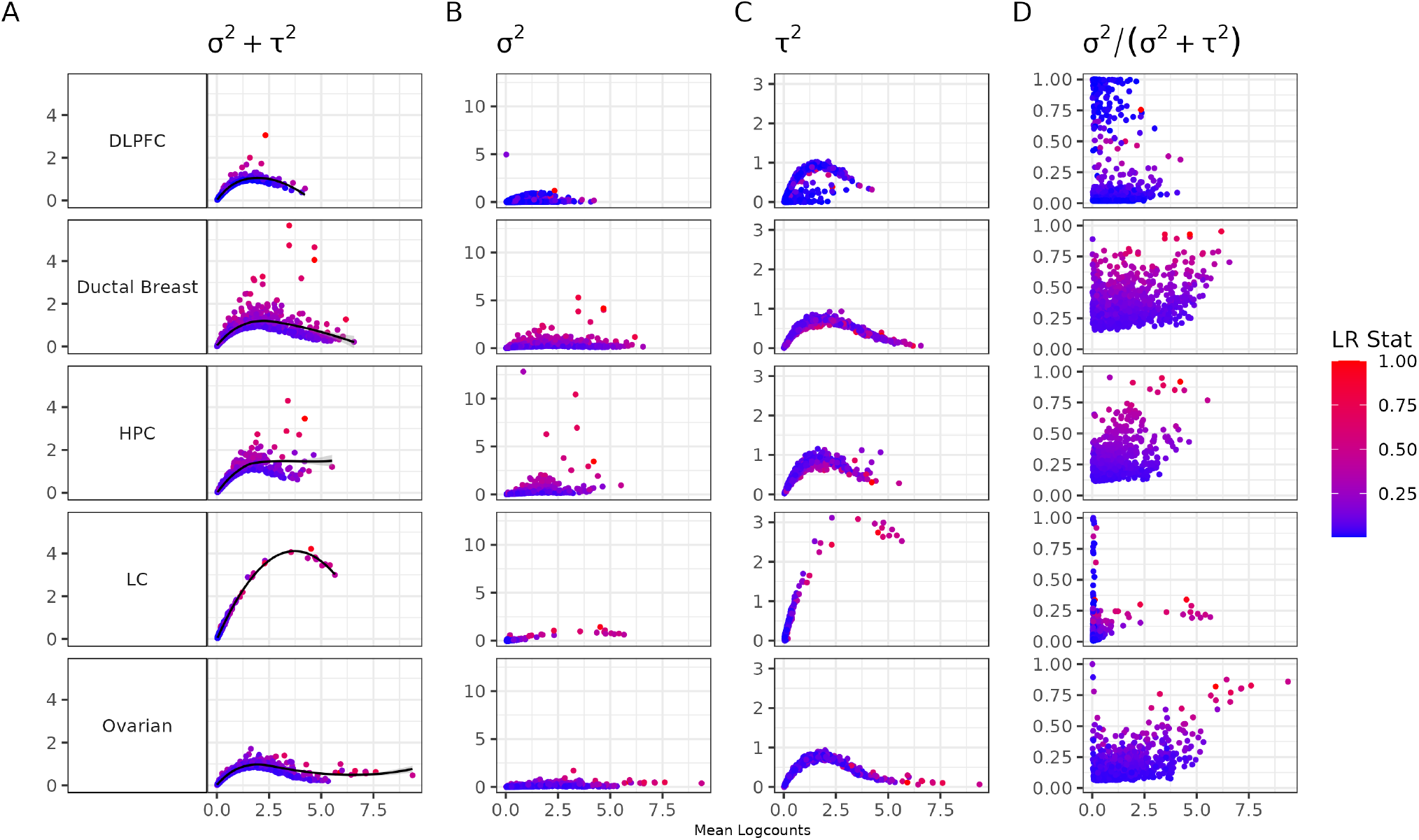
Mean-variance relationship exists in spatially resolved transcriptomics. Using data from different human tissues, in order from top to bottom: DLPFC [15], Ductal Breast cancer [39], HPC [16], LC [42], and Ovarian cancer [43], we quantified the mean-variance relationship. Each point is a gene colored by the likelihood ratio statistic for a test that compares the fitted model against a classical linear model for the spatial component of variance using a NNGP [9]. The likelihood ratio statistics (LR Stat) are scaled by the maximum likelihood ratio statistic for each dataset in order to have more uniform visualization. The x-axis is mean logcounts and the y-axes represent different components of variance, in order from left to right: total variance *σ*^2^ + *τ* ^2^, spatial variance *σ*^2^, nonspatial variance *τ* ^2^, and proportion of spatial variance *σ*^2^*/*(*σ*^2^ + *τ* ^2^).

Next, we further investigated one of these tissues (DLPFC) to ask if the mean-variance relationship was due to differences in the spatial domains of the tissue. The six layers in the human neocortex are transcriptionally quite different from one another [15], so we wanted to show that the mean-variance relationship still exists when stratifying by layers. In order to control for differences in layer domains, the DLPFC data was first separated into Layers I-VI, and white matter and then the mean logcounts were plotted against the components of variance for each layer in the brain. However, we found that the mean-variance relationship was still observed within the different biological domains (**Figure S3**).

### 3.2 The mean-rank relationship exists in other SVG detection methods

Having established that the mean-variance relationship exists in SRT data across different tissues as measured by Gaussian processes in `nnSVG`, we next explored the mean-rank relationship as an extension of the mean-variance relationship. Other SVG detection methods do not separate out the total variance into spatial and nonspatial variance components, so we examine the mean-variance relationship using this proxy.

We examined the mean-rank relationship from several popular SVG detection methods on the DLPFC, Ovarian cancer, and Lobular Breast cancer datasets (**Figure 3**). The ranks were calculated for each SVG method (described in Section 2.5). We found that for almost every method, there is a clear relationship between the mean and the rank. Stated another way, the SVG detection methods that we evaluated rank and prioritize genes as SVGs, which is related to the overall mean expression. Because the overall mean expression is likely a technical artifact, we would expect that there should be genes that are highly ranked as SVGs within each mean-level decile. However, what we found is that the mean-variance relationship biases genes towards the higher mean expression deciles. The extreme bias observed in SPARK-X is also noted in a recent benchmarking paper [48]. These are state of the art methods that perform well in recent benchmarking papers [14, 48, 49], yet they are sorely affected by the mean-variance bias.

**Figure 3:**
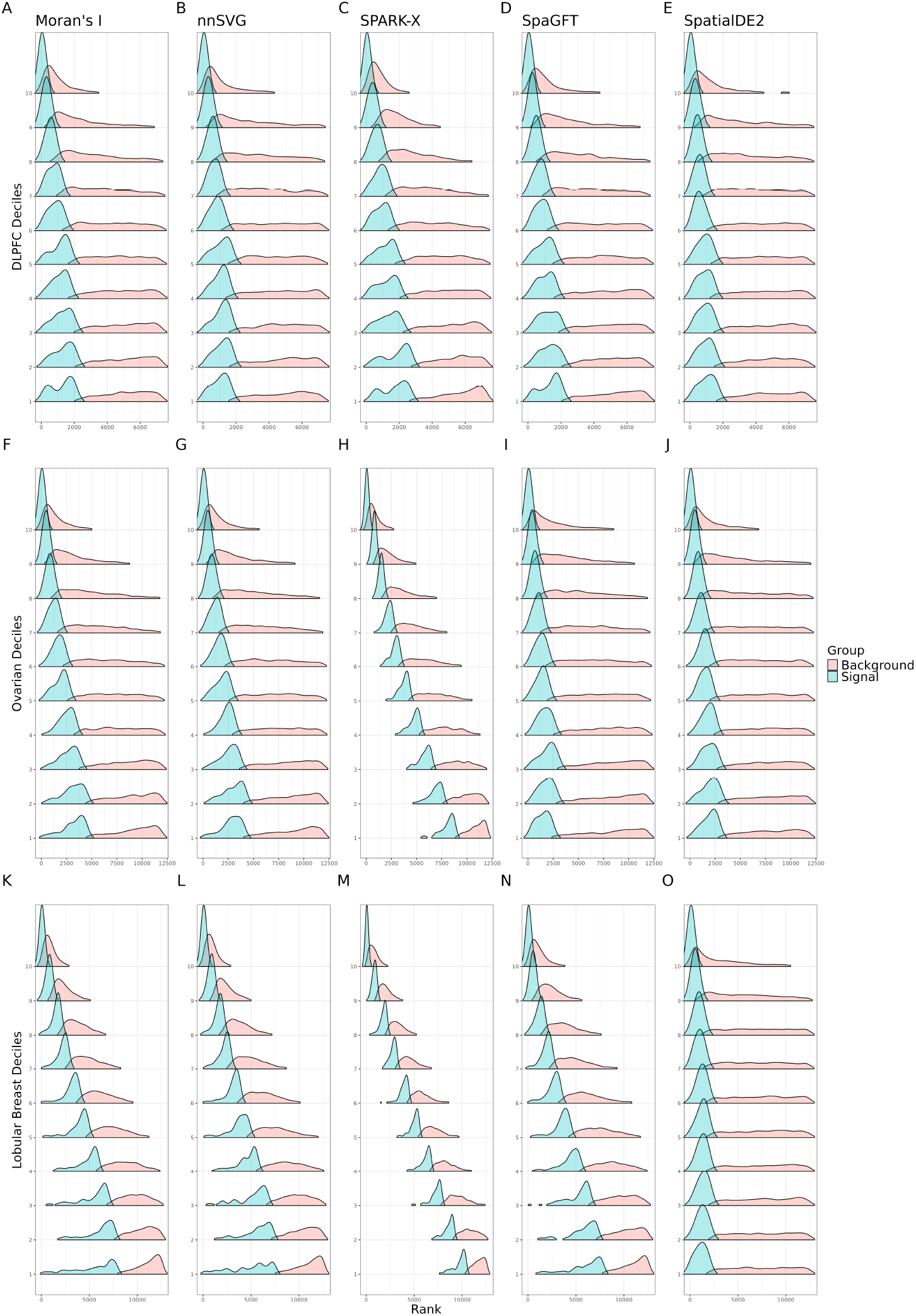
Mean-rank relationship exists in spatial transcriptomics data. Using three datasets, in order from top to bottom (DLPFC [15], Ovarian cancer [39], and Lobular Breast cancer [40]), we quantified the mean-rank relationship. The genes were binned into deciles based on mean logcounts. Decile 1 contains the lowest mean expression values. The x-axis represents the rank. Within each decile, the density of the top 10% ranks is plotted as the signal in blue, while the density of the remaining ranks is plotted as the background in orange. Each subfigure shows the mean-rank relationship that persists after applying each method, from left to right: Moran’s I [47], nnSVG [9], SPARK-X [11], SpaGFT [45], and SpatialDE2 [46].

### 3.3 Simulation: Weighted Spatially Variable Gene Evaluation

To address the mean-variance and mean-rank relationships, we began with simulation studies to evaluate the performance of *spoon* under different scenarios. Using simulated raw gene expression counts following a Poisson distribution (**Section 2.4.2**) with a fixed lengthscale (*l*=100), we ranked SVGs using nnSVG [9] without weights and with weights estimated via *spoon*. We found a strong mean-rank relationship using the unweighted SVGs (**Figure 4A**) compared to the weighted SVGs using *spoon* (**Figure 4B**). Stated differently, using observational- and gene-level weights, we can identify highly ranked SVGs even in lower deciles, demonstrating that *spoon* effectively addresses the mean-variance relationship.

**Figure 4.**
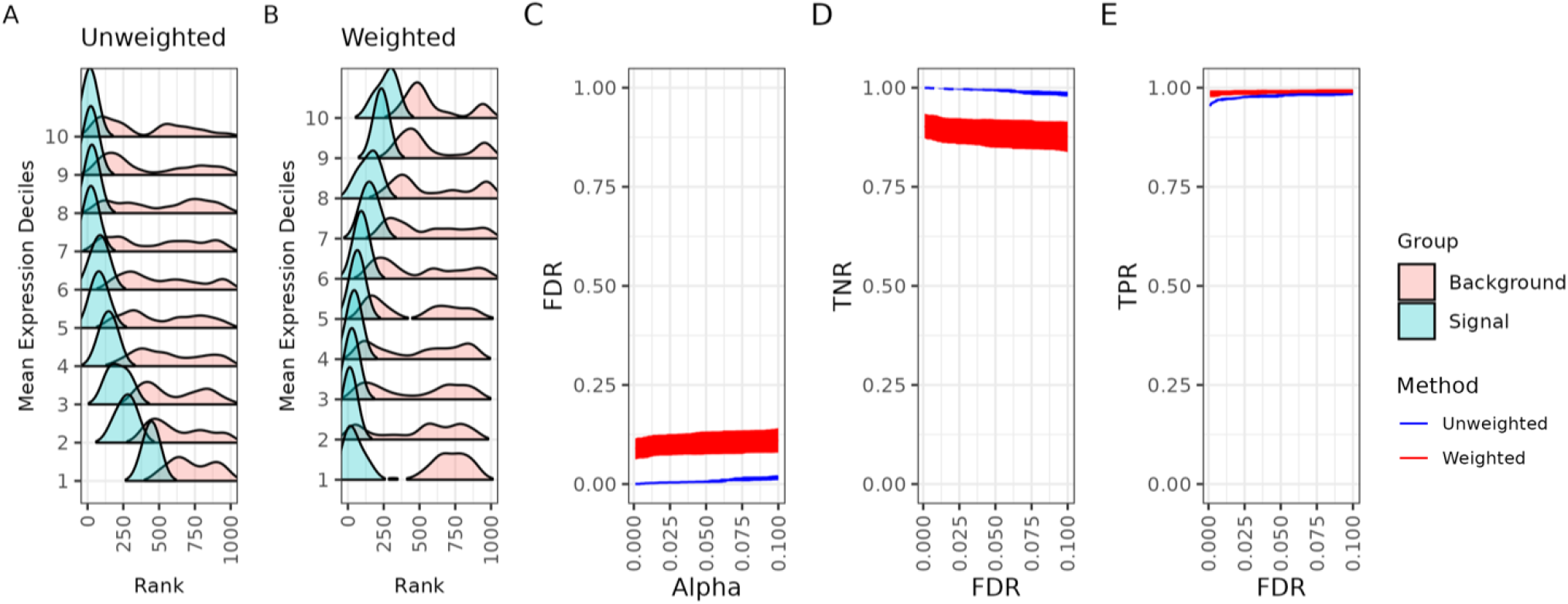
*Spoon* removes the mean-variance relationship when detecting spatially variable genes. This dataset consists of 1,000 simulated genes across 968 spots using a lengthscale of 100. Separately for unweighted and weighted methods, the genes were binned into deciles based on mean logcounts. Decile 1 contains the lowest mean expression values. Ridge plots for the (**A**) unweighted ranks and (**B**) weighted ranks are shown. Within each decile (*y*-axis), the density of the top 10% of ranks is plotted as the signal, while the density of the remaining ranks is plotted as the background. (**C**) False discovery rate (FDR) as a function of Type I error (*α*). As a function of FDR, we show the (**D**) true negative rate (TNR) and (**E**) true positive rate (TPR). The red represents weighted `nnSVG` and the blue represents unweighted `nnSVG`. These plots represent the average performance across five iterations of the same simulation, each with unique random seeds.

We also explored the false discovery rate (FDR) (**Figure 4C**), true negative rate (TNR) (**Figure 4D**), and true positive rate (TPR) (**Figure 4E**). The red represents weighted `nnSVG` and the blue represents unweighted `nnSVG`. These plots represent the average of each respective rate over five iterations of the same simulation with unique random seeds. The FDR and TNR are similar between the unweighted and weighted methods, with a slight increase in performance observed in the unweighted method. The TPR, however, is very similar for both methods. Finally, we considered other lengthscale values and found that the mean-variance relationship is improved for all values tested (**Figure S4**). We found that the weights from *spoon* improve the TPR for smaller lengthscale values, and there are diminishing returns regarding the convergence of the TPR for both the weighted method and unweighted methods at larger lengthscale values.

### 3.4 Real Data: Weighted Spatially Variable Gene Evaluation

Next, we evaluated the downstream impact of incorporating weights from *spoon* into SVG detection methods. Here, we aimed to demonstrate the impact of our method on recovering lowly expressed genes that become highly ranked in real biological datasets. We defined small mean gene expression genes as those with means less than the 25th percentile in the dataset. Within the set of small mean gene expression, we identified genes that were in the lowest 10% of ranks before weighting and then increased to the highest 10% of ranks after weighting. In the Ovarian cancer dataset, there are 7 genes that met this criterion. Out of these 7 genes, *TUFT1* and *DDX39B* are known to be implicated in ovarian cancer [50, 51]. These potentially important SVGs were ignored due to their low expression levels and our weighting algorithm can recapture them. Similar analyses were performed for the other three cancer datasets (**Figure 5**). The gene lists can be found in the supplemental materials.

**Figure 5.**
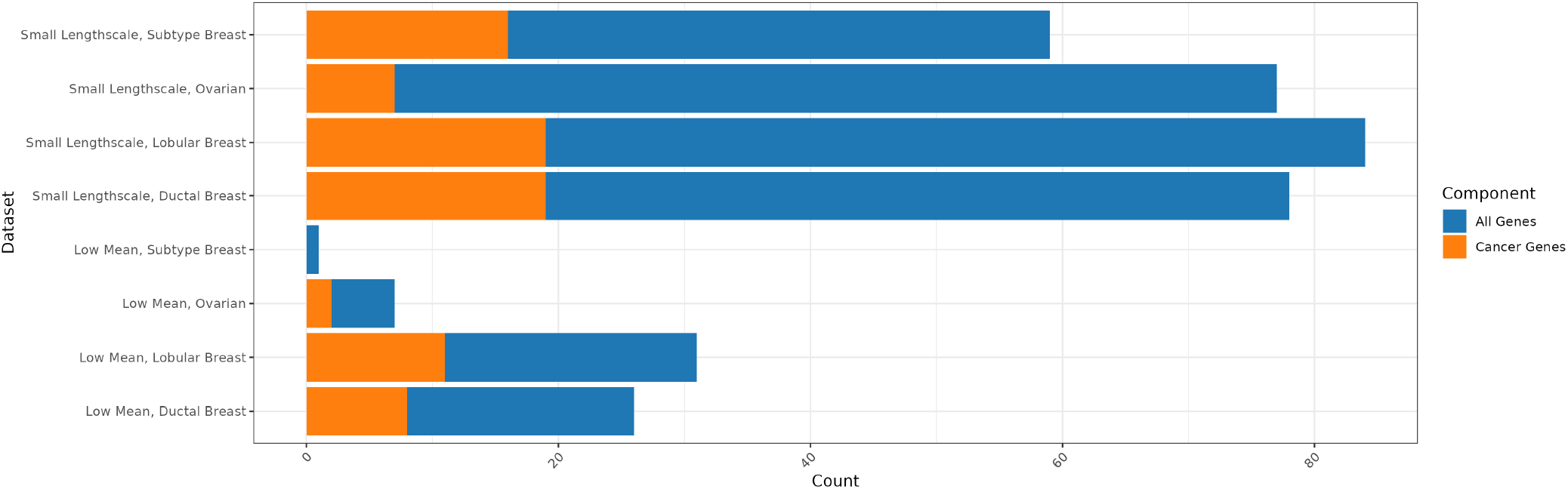
*Spoon* helps to detect SVGs associated with cancer that are lowly expressed. We used four datasets to evaluate the detection of cancer-related genes: Subtype Breast cancer [41], Ovarian cancer [43], Lobular Breast cancer [40], and Ductal Breast cancer [39]. Each bar contains the intersection of the set of genes of interest with genes within the set associated with cancer. For the first four rows, we defined low mean genes as those with means less than the 25th percentile in the dataset. Within the set of low mean genes, we found genes that were in the lowest 10% of ranks before weighting and then increased to the highest 10% of ranks after weighting. This is the set of genes of interest. The intersection in orange is the number of low mean and higher ranked genes that were found to be associated with the cancer of the dataset. For the last four rows, we defined small lengthscale genes as those with lengthscales between 40 to 90. Within the set of small lengthscale genes, we found genes that were ranked higher after weighting. This is the set of genes of interest. The intersection in orange shows the number of small lengthscale genes that were ranked higher and found to be associated with the cancer type of the dataset.

Then, we explored the improvement in the small lengthscale set of genes. We defined small length-scale genes as those with lengthscale values between 40 to 90. Within the set of small lengthscale genes, we found genes that were ranked higher after weighting. We also derived the “null distribution” — the underlying total SVGs for each dataset as a point of reference for the proportion of small lengthscale genes that are ranked higher. We found that the differing proportions of small lengthscale genes that become higher ranked after weighting is appropriate based on the “null distribution” of the proportion of unweighted SVGs (**Figure S5**). Again, we related the higher-ranked small lengthscale genes to the cancer type of the dataset. In the Subtype Breast dataset, 59 small lengthscale genes were higher ranked after weighting, with 16 of these genes implicated in breast cancer. Full results are presented in **Figure 5** and gene lists are in supplemental materials.

## 4 Discussion

In our work, we systematically demonstrate the mean-variance and the mean-rank relationships exist in spatially resolved transcriptomics data. Furthermore, we show this is not limited to just one SVG detection method. If researchers fail to adjust for this bias in spatial transcriptomics data, this can lead to false positives and inaccurate rankings of SVGs due to the violation of the homoskedasticity assumption. Here, we show that our method *spoon* is able to correct for this bias. Specifically, our approach uses Empirical Bayes techniques to generate weights for downstream analyses to remove the mean-variance relationship, leading to a more informative set of SVGs.

In a recent benchmark evaluation of SVG detection methods, the authors Chen et al. [48] noted a similar bias. NoVaTeST was recently proposed as a method to identify SVGs allowing noise variance to vary with spatial locations [52]. This method aims to identify genes that have location-dependent noise variance in SRT data, or genes that have statistically significant heteroskedasticity. This noise variation can be due to technical noise from the mean-variance relationship, variation due to sequencing processes, or underlying biological differences, making it difficult to parse out the mean-variance relationship. Additionally, further analysis of the genes detected by NoVaTest showed that some genes are likely affected by the mean-variance relationship, and the authors suggest using a strong variance-stabilizing transformation.

We recognize there are limitations to our project and aim to address these in future work. Primarily, simulation studies for spatial transcriptomics data are difficult to design and execute due to numerical instability and limitations of parameterization. There is no clear consensus on the definition of an SVG, so we chose to simulate overall SVGs, defined in Yan et al. [53] as genes that exhibit non-random spatial patterns. To our knowledge, we are not aware of methods to simulate SVGs that include the mean-variance bias. In future work, we aim to refine spatial transcriptomics simulation study design to incorporate the mean-variance relationship and have more flexibility with various parameters, such as mean gene expression, degree of spatial variation, expression strength, and varying effect sizes in the same simulated dataset. We found that our method is most powerful for small lengthscale genes, and we hope to better understand medium and large lengthscale genes in future work as well.

In sum, we provide evidence for the mean-variance and mean-rank relationship in SRT data and show that our method *spoon* can mitigate these biases. We offer the software as an easily installable R/Bioconductor package that interfaces with `SpatialExperiment` to make this method broadly accessible to researchers.

## 5 Acknowledgments

We thank members of the Hansen-Hicks lab group and our collaborators at the Lieber Institute for Brain Development for their input and feedback on this project. We also thank maintainers of the Joint High Performance Computing Exchange (JHPCE) computing cluster at Johns Hopkins Bloomberg School of Public Health for computing resources.

## Author Contributions

- **Kinnary Shah**: Conceptualization; Data curation; Formal analysis; Investigation; Methodology; Software; Validation; Visualization; Writing - original draft; Writing - review & editing
- **Boyi Guo**: Investigation; Methodology; Software; Visualization; Writing - review & editing
- **Stephanie C. Hicks**: Conceptualization; Formal analysis; Funding acquisition; Investigation; Methodology; Project administration; Resources; Software; Supervision; Validation; Visualization; Writing - review & editing

## Supplementary Materials

**Figure S1:**
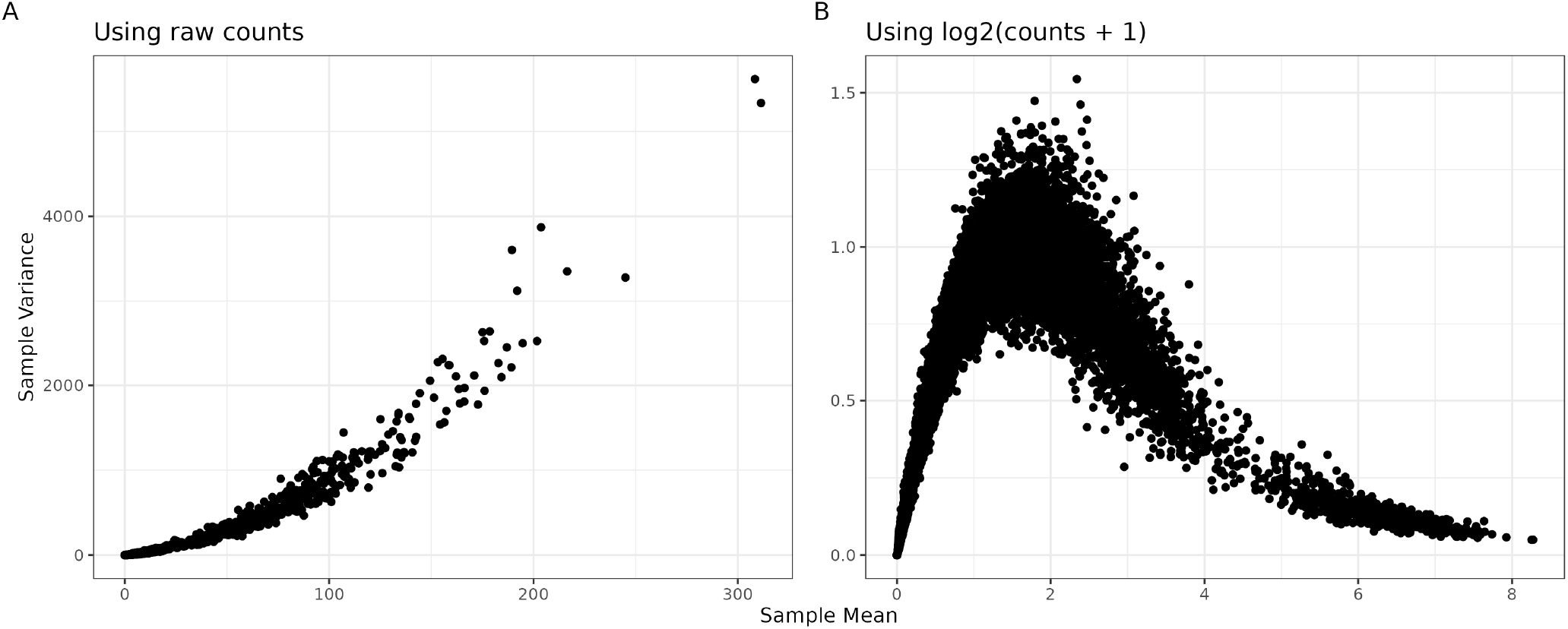
Visualizing the mean-variance relationship on different scales. The mean-variance relationship exists with or without a log-transformation. Gene expression counts were simulated using the `splatter` R/Bioconductor package [54] for *G*=10,000 genes and *N* =100 observations (or cells) under a Gamma-Poisson model. Each point represents one gene. Both representations illustrate the mean-variance relationship where the *x*-axis is the sample mean and *y*-axis is the sample variance using either (**A**) the raw counts or (**B**) the log_2_-transformed counts with a pseudocount of 1 (or log_2_(counts+1)). Here, the log-transformation overcorrects for the mean-variance relationship for the larger counts.

**Figure S2:**
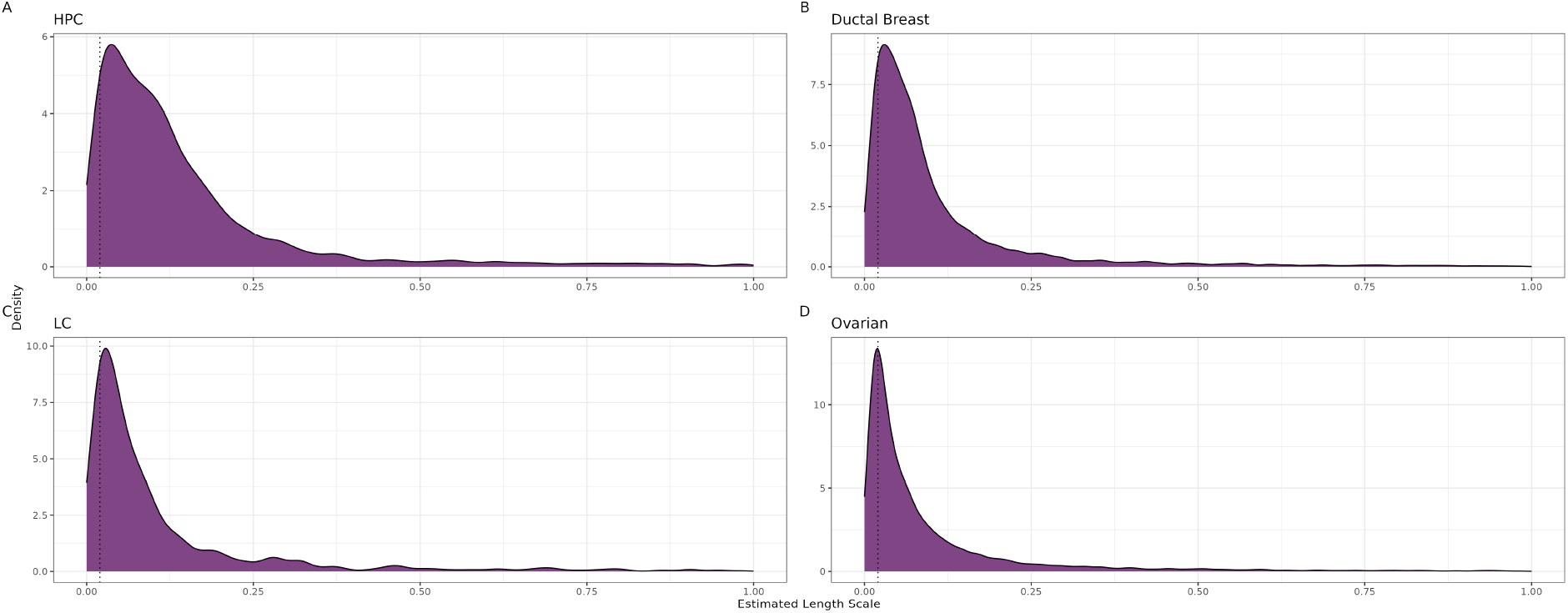
Real data estimated lengthscale distributions using nnSVG. This figure shows the estimated lengthscale distributions for four real datasets (**A**) HPC [16], (**B**) Ductal Breast cancer [39], (**C**) LC [42], and (**D**) Ovarian cancer [43]. For each dataset, `nnSVG` was used to calculate the estimated lengthscale value for each gene and the distribution of values between 0 and 1 is plotted. The dotted line highlights the lengthscale value used in the primary simulation evaluations.

**Figure S3:**
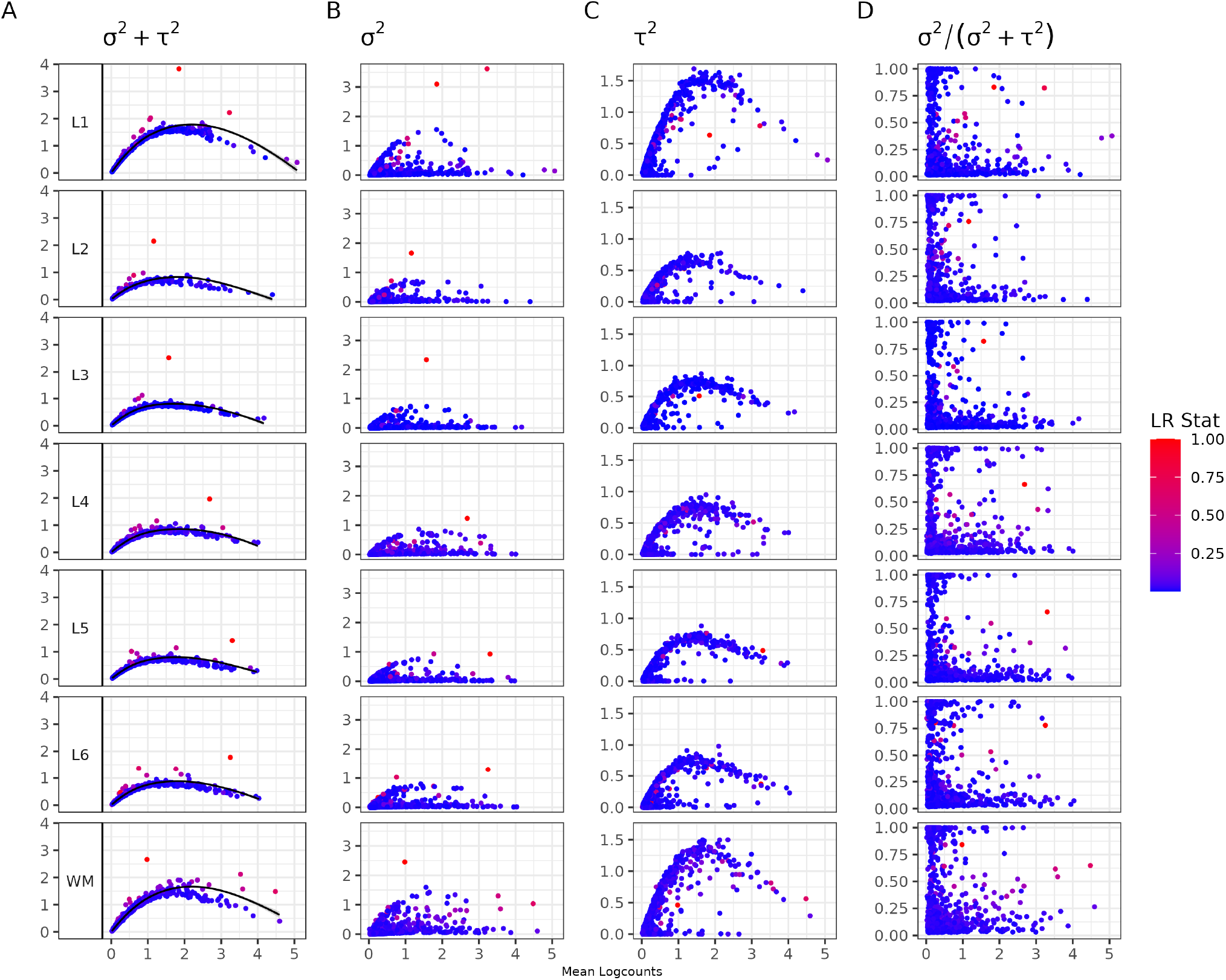
Mean-variance relationship after conditioning out biological variance measured by Gaussian process. Each row is a cortical layer from the DLPFC dataset, in order from top to bottom: Layers I-VI, white matter (WM). Each point is a gene colored by the likelihood ratio statistic (LR Stat) for a test comparing the fitted model against a classical linear model for the spatial component of variance. The likelihood ratio statistics are scaled by the maximum likelihood ratio statistic for each layer in order to have more uniform visualization. The x-axis represents mean logcounts and the y-axes represent different components of variance, in order from left to right: total variance *σ*^2^ + *τ* ^2^, spatial variance *σ*^2^, nonspatial variance *τ* ^2^, and proportion of spatial variance *σ*^2^*/*(*σ*^2^ + *τ* ^2^).

**Figure S4:**
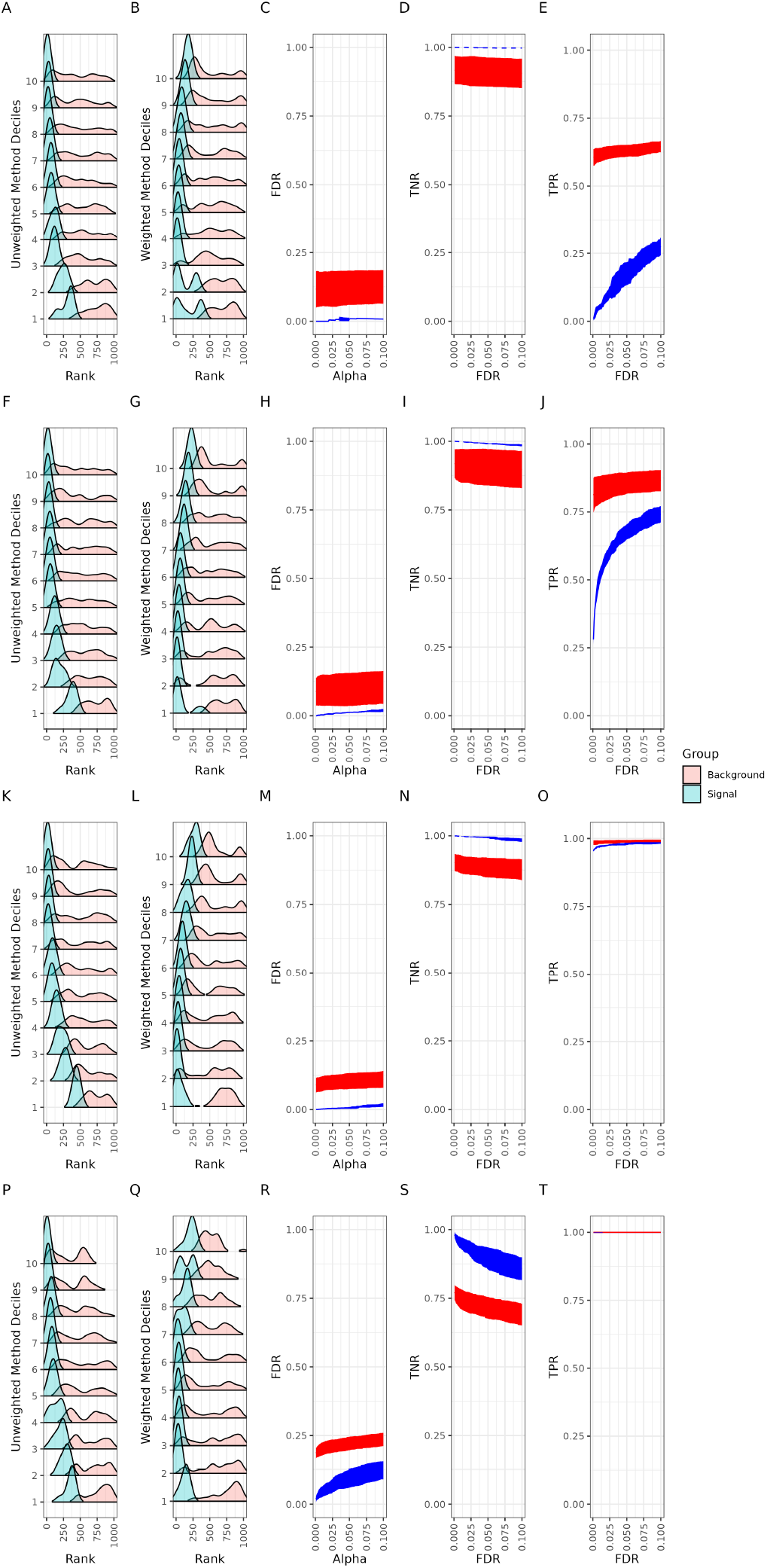
Removing the mean-variance relationship with expanded lengthscale metrics. This dataset contains 1,000 simulated genes across 968 spots. Each row represents a simulation setting with unique lengthscales, in order from top to bottom: 50, 60, 100, 500. Separately for unweighted and weighted methods, the genes were binned into deciles based on mean logcounts. Decile 1 is the lowest mean expression values. The first column of plots is unweighted ranks and the second column of plots is weighted ranks. Within each decile, the density of the top 10% ranks is plotted as the signal and the density of the remaining ranks is plotted as the background. The final three columns show the false discovery rate (FDR), true negative rate (TNR), and true positive rate (TPR). The red represents weighted nnSVG and the blue represents unweighted nnSVG. These plots represent the average of each respective rate over five iterations of the same simulation with unique random seeds.

**Figure S5:**
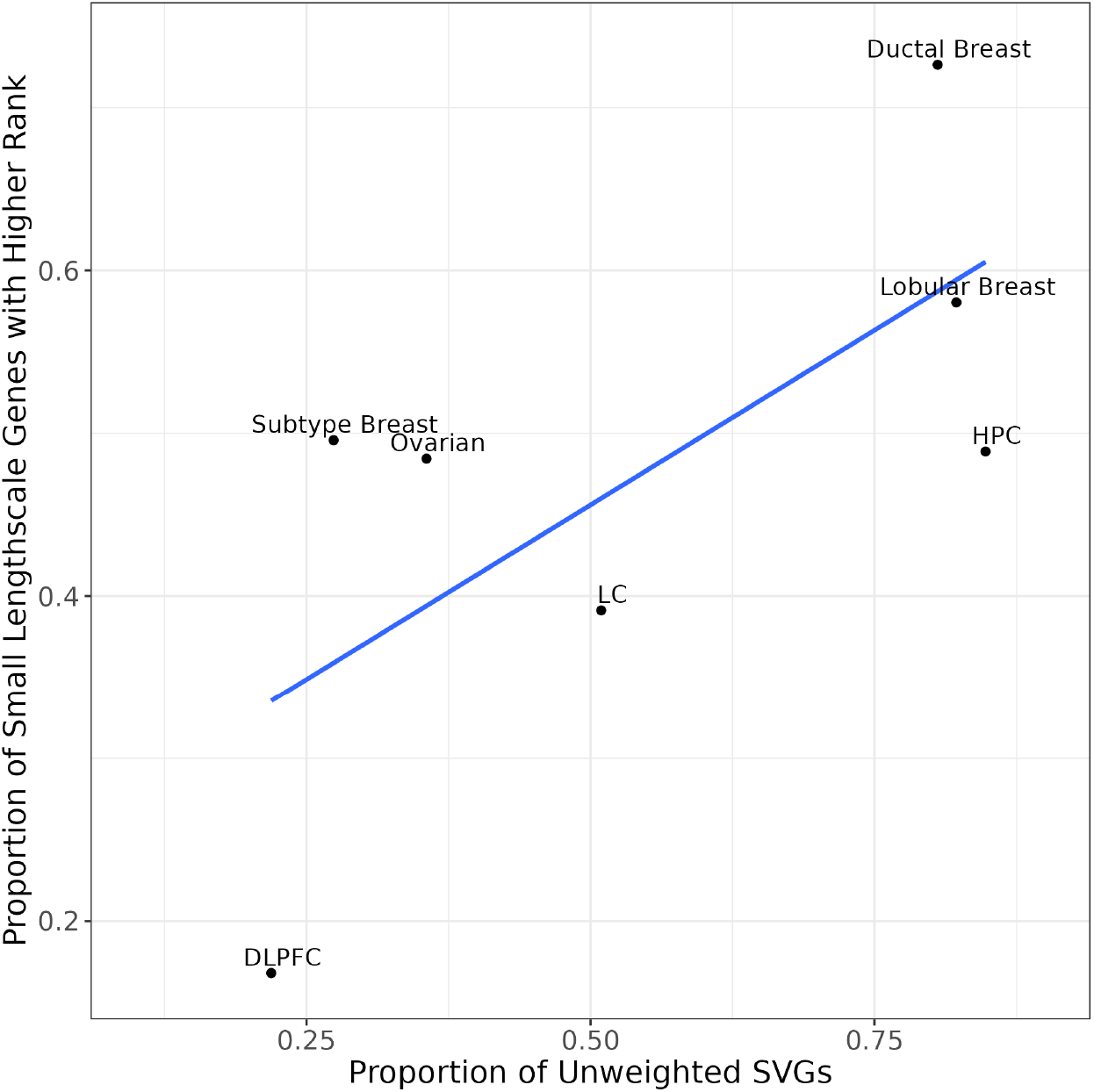
Ranking small lengthscale genes after weighting. Each point is a unique real dataset analyzed with 10x Genomics Visium. The x-axis is the proportion of SVGs from running unweighted `nnSVG` on the dataset. The y-axis is the proportion of genes with a small lengthscale (40-90) that are higher ranked in weighted `nnSVG` compared to unweighted `nnSVG`.

**Table S1:**
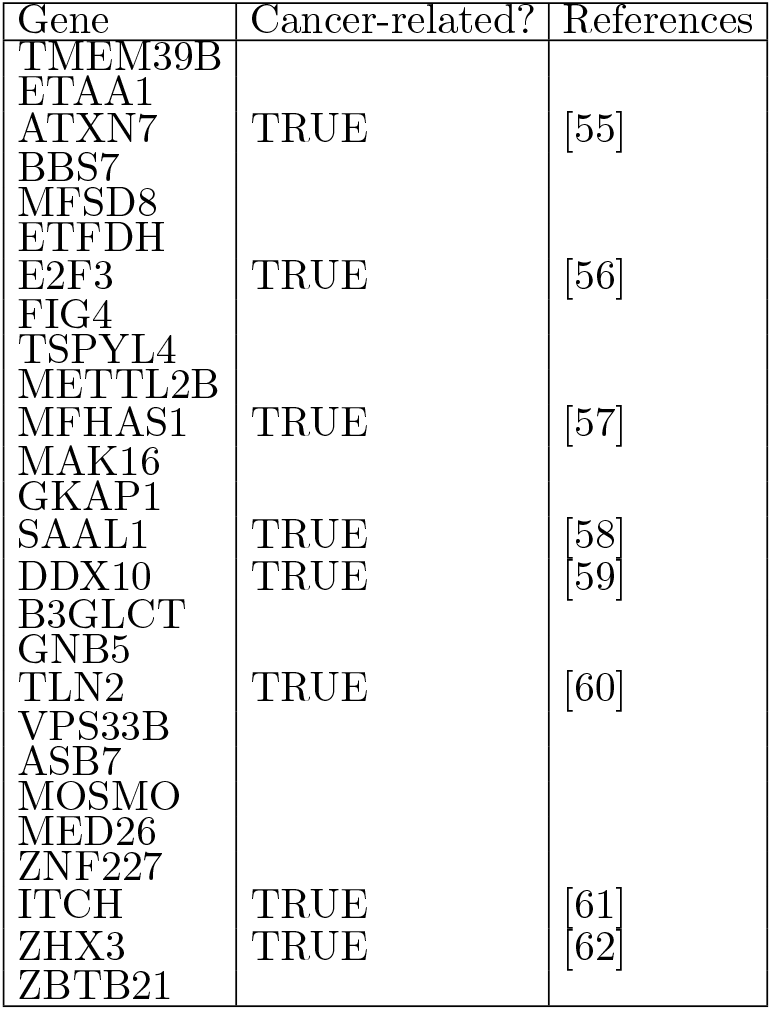
Low Mean, Ductal Breast Genes Relation to Cancer. This table shows all genes with means less than the 25th percentile in the Ductal Breast [39] dataset which were in the lowest 10% of ranks before weighting and then increased to the highest 10% of ranks after weighting. The second column indicates if the gene is related to Breast cancer, with the corresponding reference in the third column.

**Table S2:**
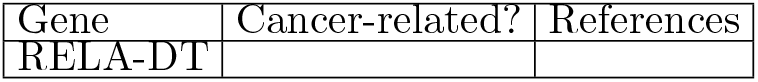
Low Mean, Subtype Breast Genes Relation to Cancer. This table shows all genes with means less than the 25th percentile in the Subtype Breast [41] dataset which were in the lowest 10% of ranks before weighting and then increased to the highest 10% of ranks after weighting. The second column indicates if the gene is related to Breast cancer, with the corresponding reference in the third column.

| Gene | Cancer-related? | References |
| --- | --- | --- |
| RELA-DT |  |  |

**Table S3:**
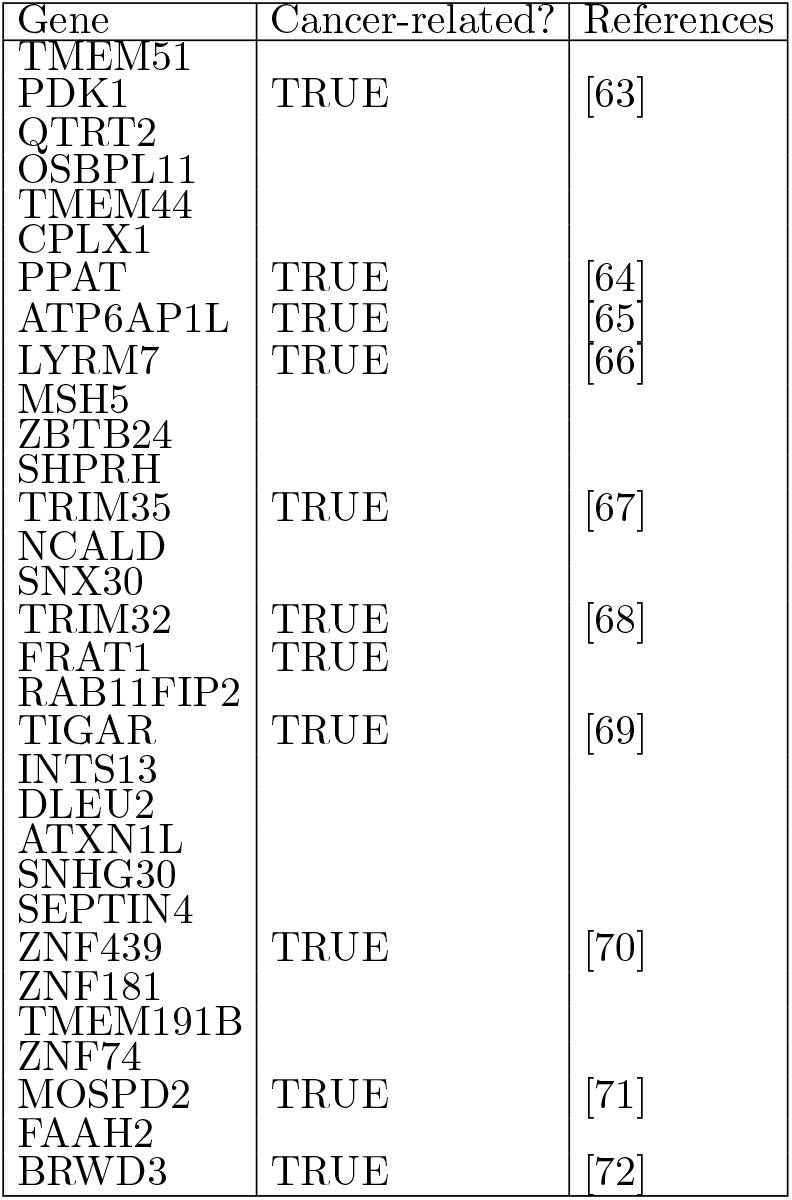
Low Mean, Lobular Breast Genes Relation to Cancer. This table shows all genes with means less than the 25th percentile in the Lobular Breast [40] dataset which were in the lowest 10% of ranks before weighting and then increased to the highest 10% of ranks after weighting. The second column indicates if the gene is related to Breast cancer, with the corresponding reference in the third column.

**Table S4:**
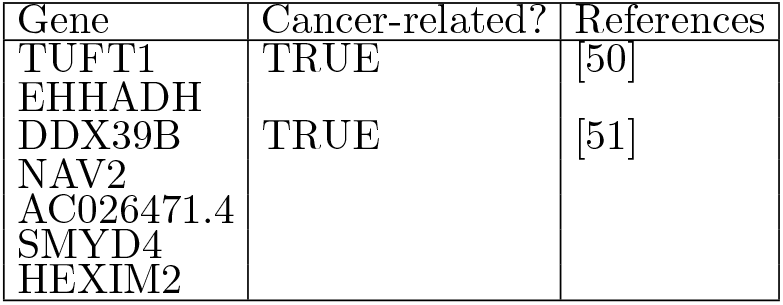
Low Mean, Ovarian Genes Relation to Cancer. This table shows all genes with means less than the 25th percentile in the Ovarian [43] dataset which were in the lowest 10% of ranks before weighting and then increased to the highest 10% of ranks after weighting. The second column indicates if the gene is related to Ovarian cancer, with the corresponding reference in the third column.

**Table S5:**
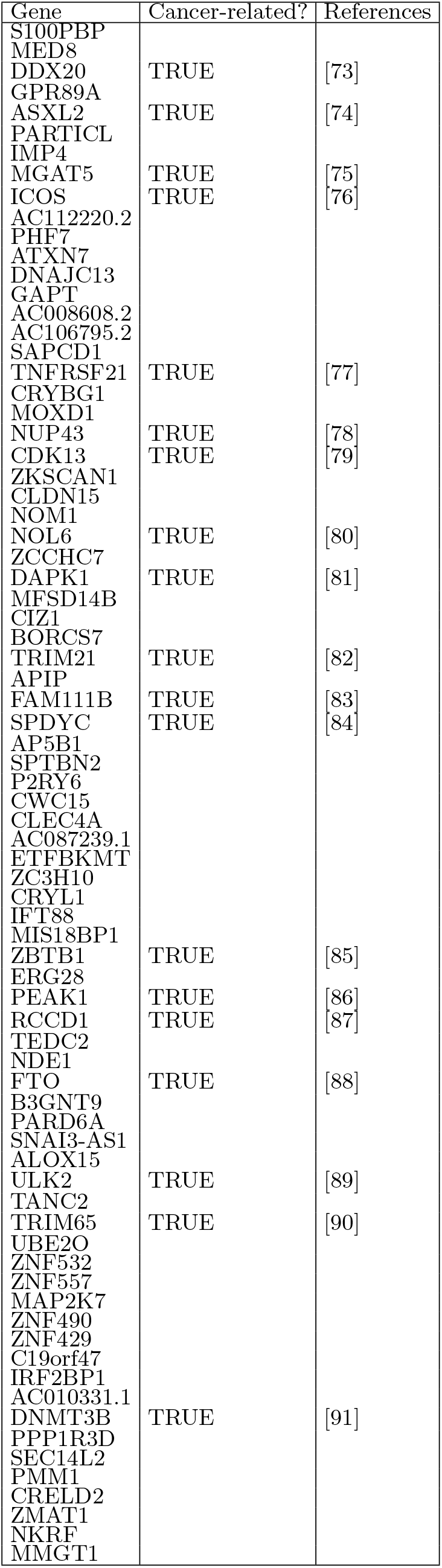
Small Lengthscale, Ductal Breast Genes Relation to Cancer. This table shows all genes with lengthscale values between 40 to 90 in the Ductal Breast [39] dataset which were ranked higher after weighting. The second column indicates if the gene is related to Breast cancer, with the corresponding reference in the third column.

**Table S6:**
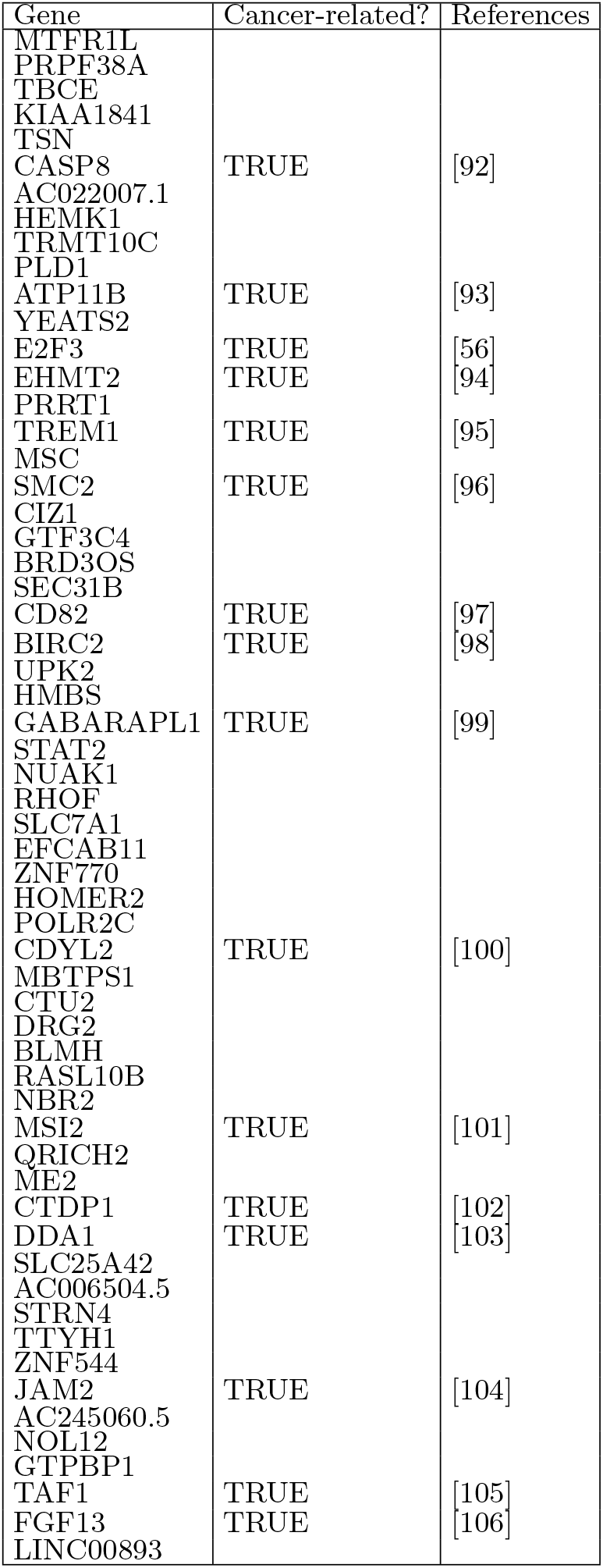
Small Lengthscale, Subtype Breast Genes Relation to Cancer. This table shows all genes with lengthscale values between 40 to 90 in the Subtype Breast [41] dataset which were ranked higher after weighting. The second column indicates if the gene is related to Breast cancer, with the corresponding reference in the third column.

**Table S7:**
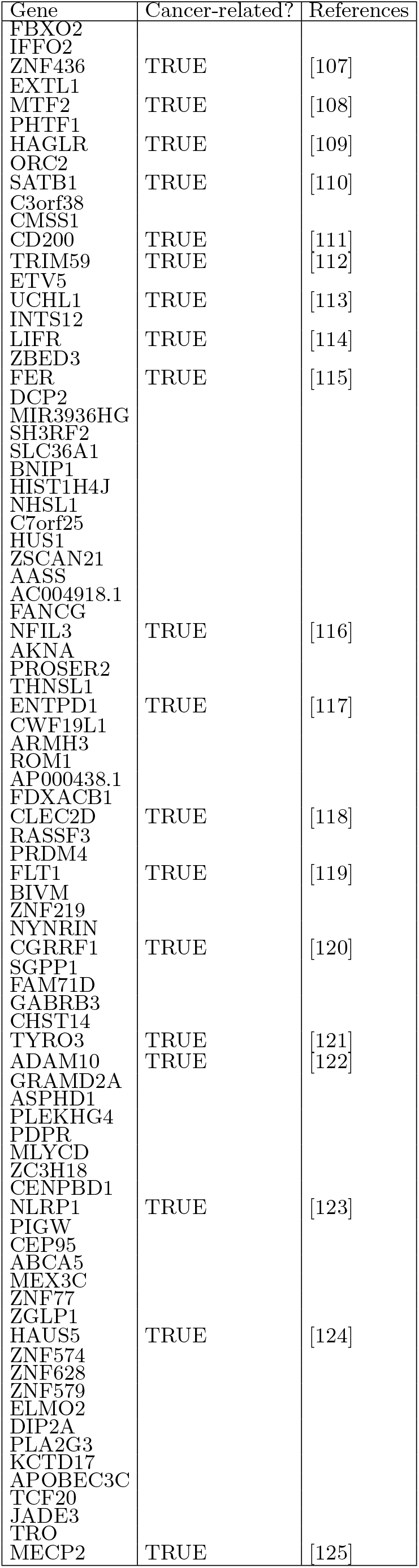
Small Lengthscale, Lobular Breast Genes Relation to Cancer. This table shows all genes with lengthscale values between 40 to 90 in the Lobular Breast [40] dataset which were ranked higher after weighting. The second column indicates if the gene is related to Breast cancer, with the corresponding reference in the third column.

**Table S8:**
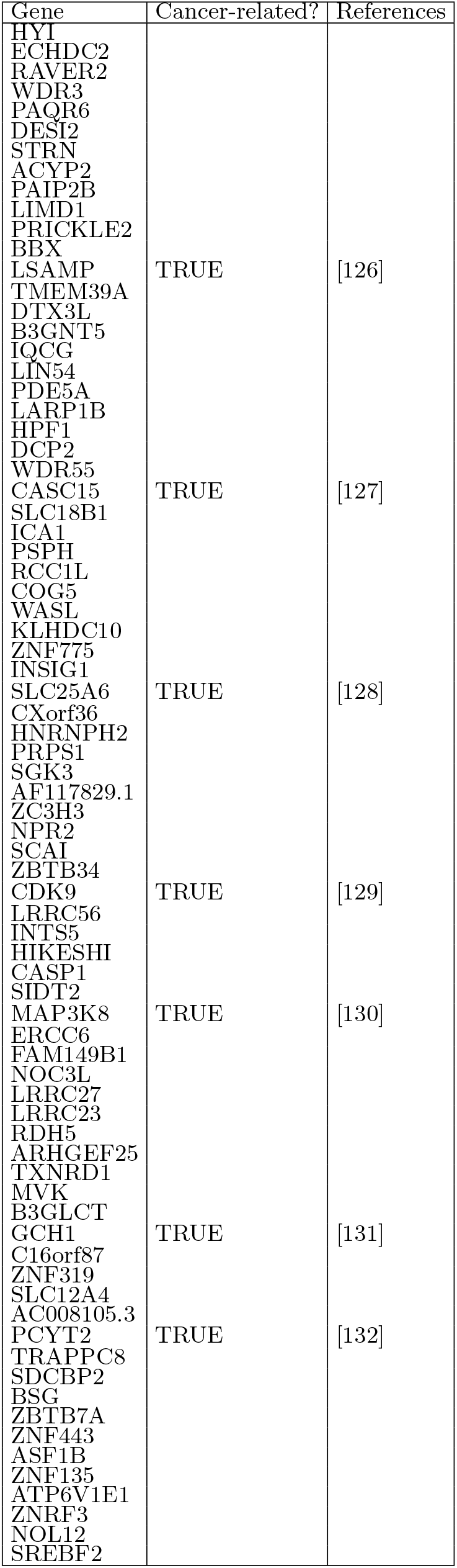
Small Lengthscale, Ovarian Genes Relation to Cancer. This table shows all genes with length-scale values between 40 to 90 in the Ovarian [43] dataset which were ranked higher after weighting. The second column indicates if the gene is related to Ovarian cancer, with the corresponding reference in the third column.

